# Genomic and regulatory basis of adaptation in Cameroonian Gudali and Simgud Cattle

**DOI:** 10.1101/2025.10.13.680102

**Authors:** Youchahou Poutougnigni Matenchi, Matthew Hegarty

## Abstract

**Background:** Gudali, a West and Central African shorthorn zebu renowned for its dual-purpose potential, is a key genetic resource in regional livestock production. It has recently been used in major crossbreeding programs, notably with Italian Simmental, to produce the Simgud hybrid. These initiatives aim to combine the exceptional adaptive traits of Gudali with the superior productive performance of Simmental. However, the genomic impact of such crossbreeding on both adaptation and performance remains poorly understood. In this study, we investigated candidate signatures of selection and their associations with quantitative trait loci (QTL) and functionally important genes in the genomes of Gudali and Simgud. Our findings provide insights to guide reasoned, targeted breeding strategies that enhance productivity in tropical environments while preserving adaptive potential.

**Results:** From a dataset of 539 Gudali and 139 Simgud genotyped with the GeneSeek GGP+ Bovine 100K array, we performed a two-step imputation to whole genome and used the resulting dataset to detect candidate selection signatures using Tajima’s D, the integrated haplotype score (iHS), the fixation index (F_ST_), and the cross-population extended haplotype homozygosity (XP-EHH). Combining the identified regions under selection, together with gene expression and quantitative trait loci (QTL) databases, we further investigated the genomic targets of natural and artificial selection to identify functional candidate genes underlying adaptation mechanisms. In general, the regions under selection were associated mainly with immunity, food scarcity, thermotolerance and various production traits as important selection targets. For instance, signals detected on BTA5 and BTA7 shared between Gudali and Simgud harbored many olfactory genes (OR2O2, OR7A94, OR7H5P) and taste receptors in Simgud (TAS2R42 and TAS2R46) essential to detect forages and predators in grazing lands. Analyzing 27 tissues, we found that the genes within the regions under selection were mostly enriched for those overexpressed in testis, lung, kidney and hypothalamus.

**Conclusion:** By integrating signatures of selection with information from QTL and gene expression, we identified four genes whose relevance was supported not only by selection signals but by additional functional evidence. For instance, GAB2 for response to trypanosome infection and EYA1 associated with heat/drought adaptation, both needed to thrive in challenging tropical environments.

## Introduction

Understanding the genetic basis of adaptation and productivity in livestock is essential for improving animal breeding strategies, particularly in environments characterized by diverse ecological pressures^1^. In sub-Saharan Africa, local cattle breeds such as the indigenous Gudali have evolved under natural selection for traits such as immune response, disease resistance, heat tolerance, and low-input productivity^2–6^. Meanwhile, crossbred populations like the Simgud (a cross between Gudali and Simmental) offer a unique opportunity to study how artificial selection and admixture^7^ influence the genomic landscape of adaptation. This study investigates genomic signatures of selection and adaptive regulatory variants in Gudali and Simgud cattle of Cameroon. We employed a combination of robust population genomic metrics - including integrated Haplotype Score (iHS), Tajima’s D, Cross Population Extended Haplotype Homozygosity (XP-EHH), and Fixation Index (F_ST_) - to detect recent and divergent selection events across the genome. To provide functional insights, we mapped the identified selection signals to known Quantitative Trait Loci (QTL) and expression Quantitative Trait Loci (eQTL), allowing us to associate genomic variation with traits of economic and adaptive relevance. Linking signatures of selection with gene expression data is suggested as a means of gaining deeper insight into trait-associations of selection^8^. Such studies have improved the understanding of adaptation mechanisms in humans and chickens^8–10^. However, because they need large sample sizes for accuracy of eQTL detection, they are cost prohibitive - especially in developing countries, where they are almost unfeasible. The cattle Genotype-Tissue-Expression atlas (CattleGTEx) - an extensive public database of tissue-specific gene expression levels, expression quantitative-trait loci (eQTL) and transcriptome-wide associations^11^ - provides a foundation for such analysis under resource limited conditions. Several studies have used this resource and revealed important trait-associations of selection signatures in Chinese Holstein^12^, American beef cattle^13^ and more recently in a limited size sample of some African breeds^14^. Here we used the CattleGTEx data on the largest genotyped population of local Gudali and its crossbreed with the Italian Simmental (Simgud), in order to understand how both natural and artificial selection can shape the genome of these populations. By integrating selective sweep detection with QTL and eQTL annotation, our findings highlight candidate genomic regions and regulatory variants likely involved in local adaptation and breed-specific selection pressures. These results contribute to a better understanding of the genetic architecture underlying adaptation in African cattle and provide genomic resources that can inform future breeding and conservation strategies.

## Results

### Within population selection signatures

#### iHS Test Statistic

Within population analyses of selection signals using the iHS statistic identified several genomic regions exhibiting elevated Z-scores. Regions containing at least one variant within the top 0.01% of the genome-wide Z-score distribution were considered putative selection signatures. A total of 16 and 26 such regions were identified in Gudali (Fig. 1) and Simgud (Fig. 2) populations, respectively and contained 9 (Gudali) and 23 (Simgud) genes, as shown in Table 1. Most of these regions were shared either partially or entirely between the two populations. However, some breed-specific signals were observed: regions on BTA9, BTA15 and BTA23 were unique to Gudali, while signals on BTA2, BTA3, BTA4, BTA14, BTA20, and BTA22 were exclusive to Simgud.

**Table 1.** Genomic regions under selection and corresponding candidate genes.

| Population | BTA | Region start | Region end | Region length | Gene start | Gene end | Gene name |
| --- | --- | --- | --- | --- | --- | --- | --- |
| Gudali | 1 | 115526583 | 115676498 | 149915 | NA | NA | NA |
|  | 1 | 115765301 | 115847195 | 81894 | NA | NA | NA |
|  | 1 | 155669221 | 155974834 | 305613 | NA | NA | NA |
|  | 5 | 57674065 | 57683913 | 9848 | 57673709 | 57674653 | OR6C8 |
|  | 6 | 4903949 | 5360037 | 456088 | 4937484 | 4937590 | 5S_rRNA |
|  | 7 | 10384261 | 10433083 | 48822 | 10416394 | 10417392 | ENSBTAG00000052896 |
|  | 7 | 10384261 | 10433083 | 48822 | 10428807 | 10442492 | ENSBTAG00000055231 |
|  | 7 | 40173732 | 40182100 | 8368 | 40178438 | 40179370 | OR2O2 |
|  | 8 | 111803097 | 111887450 | 84353 | NA | NA | NA |
|  | 9 | 71806691 | 71843460 | 36769 | NA | NA | NA |
|  | 11 | 17373447 | 17400577 | 27130 | 17395111 | 17395172 | U7 |
|  | 15 | 84342769 | 84467632 | 124863 | 84390783 | 84391700 | OR4A2I |
|  | 15 | 84342769 | 84467632 | 124863 | 84426101 | 84427018 | OR4A2C |
|  | 23 | 8038702 | 8103193 | 64491 | NA | NA | NA |
|  | 24 | 55841983 | 55903947 | 61964 | 55937973 | 55981193 | TXNL1 |
|  | 24 | 55973886 | 56011681 | 37795 | 56001377 | 56351749 | WDR7 |
|  | 29 | 424195 | 441672 | 17477 | NA | NA | NA |
| Simgud | 1 | 115769662 | 115821790 | NA | NA | NA | NA |
|  | 1 | 140060595 | 140166434 | 105839 | 140132680 | 140470932 | DSCAM |
|  | 2 | 10971939 | 11060520 | NA | NA | NA | NA |
|  | 2 | 122192786 | 122211777 | 18991 | 122183940 | 122198209 | ENSBTAG00000039994 |
|  | 3 | 13268796 | 13271876 | NA | NA | NA | NA |
|  | 3 | 13290226 | 13303347 | NA | NA | NA | NA |
|  | 3 | 13304275 | 13368735 | 64460 | 13352587 | 13381774 | ENSBTAG00000050921 |
|  | 3 | 27448612 | 27513931 | 65319 | 27474900 | 27478019 | NHLH2 |
|  | 4 | 4593325 | 4610629 | 17304 | 4582609 | 4879280 | COBL |
|  | 5 | 47759148 | 47888834 | 129686 | 47819505 | 47966760 | HMGA2 |
|  | 5 | 98474503 | 98563105 | 88602 | 98522812 | 98558643 | ENSBTAG00000048489 |
|  | 5 | 98587898 | 98597173 | 9275 | 98549494 | 98550424 | TAS2R42 |
|  | 5 | 98587898 | 98597173 | 9275 | 98588763 | 98589680 | TAS2R46 |
|  | 5 | 98587898 | 98597173 | 9275 | 98595392 | 98596309 | ENSBTAG00000001336 |
|  | 6 | 4910894 | 5420292 | 509398 | 4937484 | 4937590 | 5S_rRNA |
|  | 7 | 10287261 | 10319408 | 32147 | 10296528 | 10297538 | OR7A94 |
|  | 7 | 10381997 | 10436556 | 54559 | 10416394 | 10417392 | ENSBTAG00000052896 |
|  | 7 | 10381997 | 10436556 | 54559 | 10428807 | 10442492 | ENSBTAG00000055231 |
|  | 7 | 60586934 | 60761559 | 174625 | 60505700 | 60641090 | ABLIM3 |
|  | 7 | 60586934 | 60761559 | 174625 | 60664747 | 60740423 | AFAP1L1 |
|  | 7 | 60586934 | 60761559 | 174625 | 60743594 | 60752561 | GRPEL2 |
|  | 7 | 60586934 | 60761559 | 174625 | 60754588 | 60767542 | PCYOX1L |
|  | 7 | 13676054 | 13699248 | 23194 | 13697483 | 13698416 | OR7H5P |
|  | 8 | 104714623 | 104820170 | NA | NA | NA | NA |
|  | 20 | 14838139 | 14862179 | 24040 | 14840060 | 14973475 | RGS7BP |
|  | 20 | 40123250 | 40138918 | 15668 | 39873127 | 40265889 | ADAMTS12 |
|  | 22 | 31593548 | 31935449 | 341901 | 31617620 | 31843533 | MITF |
|  | 24 | 56169331 | 56263062 | 93731 | 56001377 | 56351749 | WDR7 |
|  | 29 | 433088 | 480656 | NA | NA | NA | NA |

**Figure 1.**
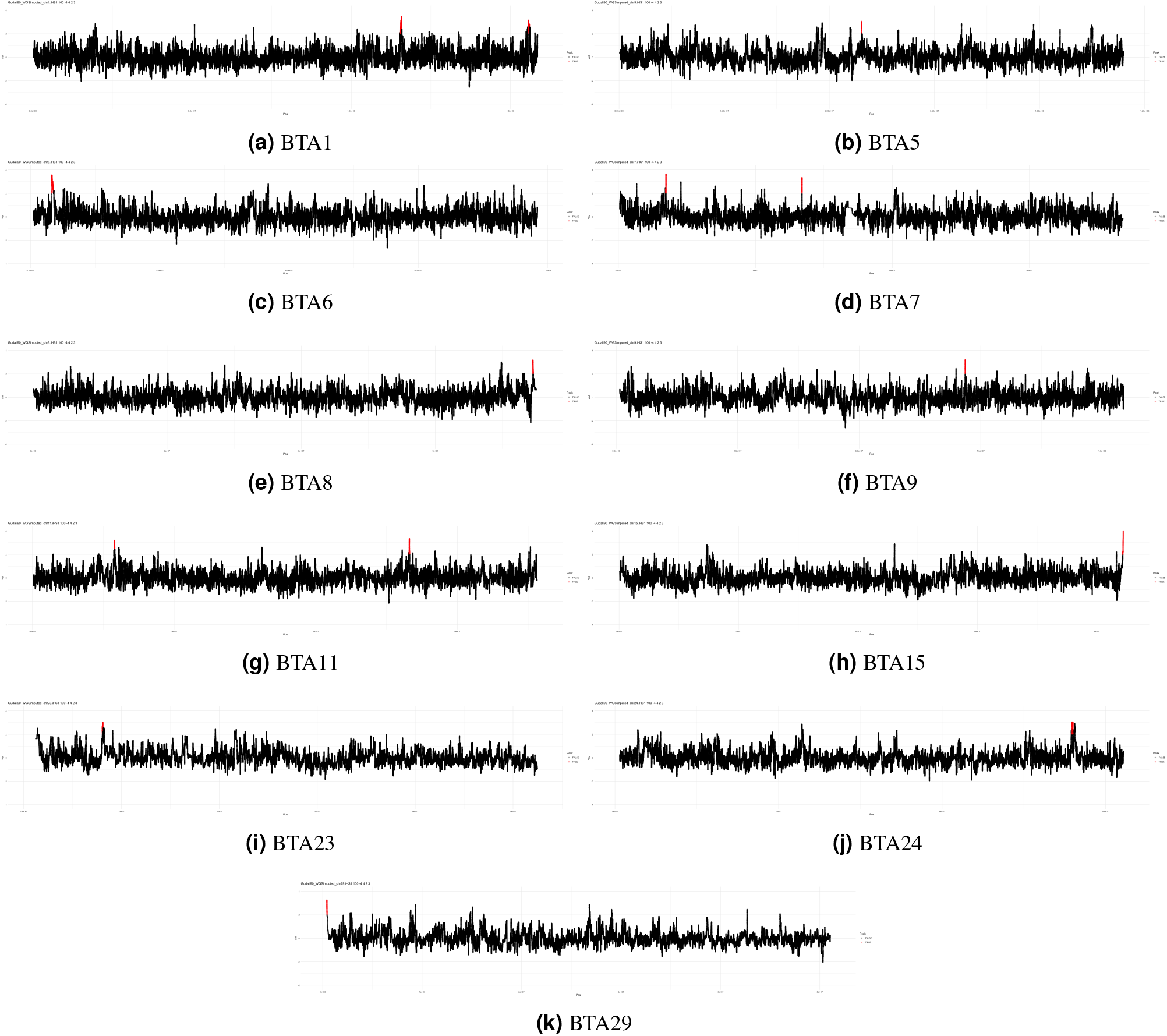
Standardized iHS values across selected chromosomes in the Gudali population.

**Figure 2.**
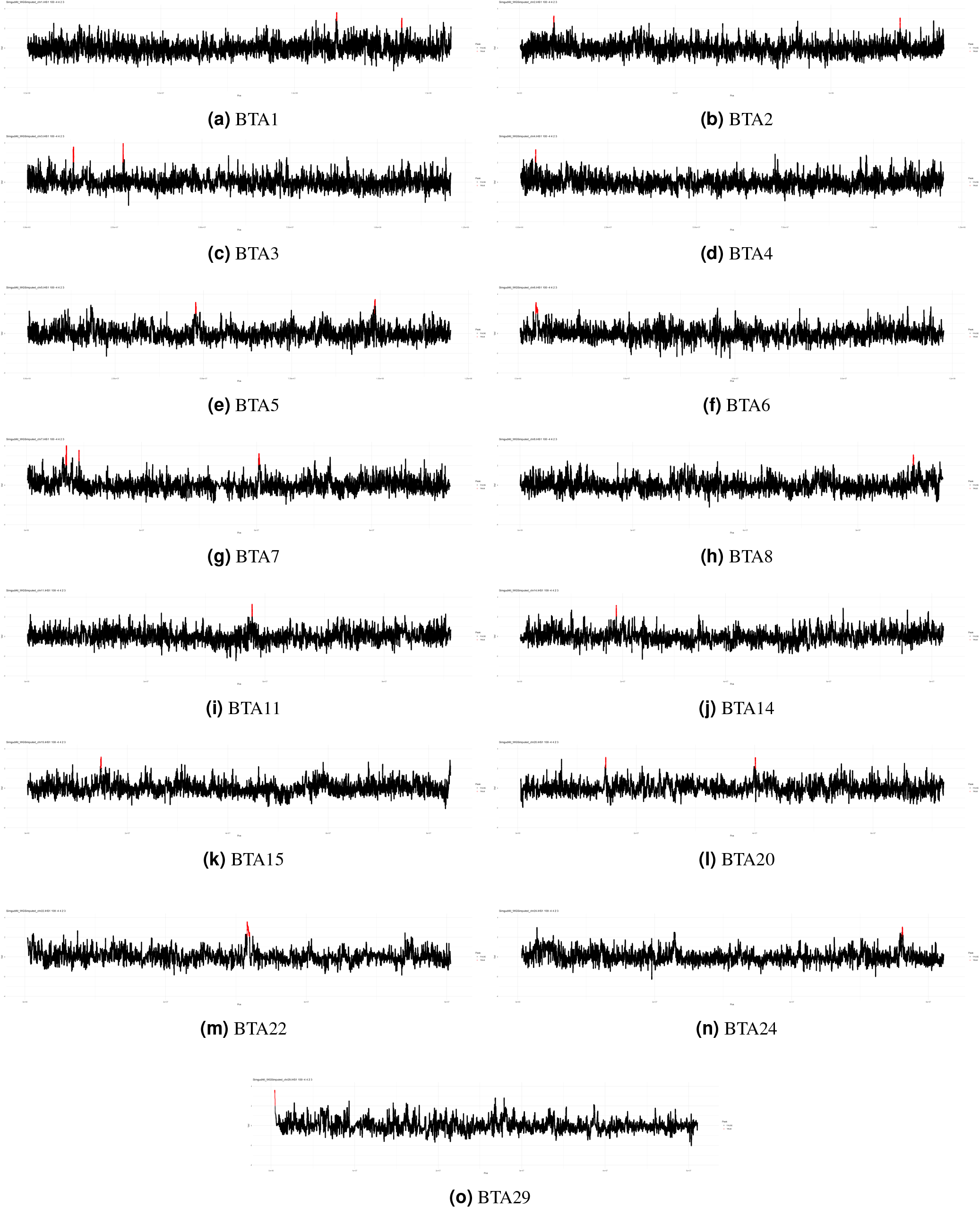
Standardized iHS values across selected chromosomes in the Simgud population.

#### Signatures of selection found in Gudali only

Two candidate regions showing signatures of selection specific to Gudali were identified on BTA23 and BTA9, but these did not contain any known genes. However, the larger region (spanning 8.04–8.10 Mb on BTA23) overlaps with a QTL associated with marbling score (QTL ID: 232813), a trait linked to meat quality. Similarly, the region between 71.81–71.84 Mb on BTA9 coincides with a QTL related to non-return rate (QTL ID: 15296), which is indicative of fertility performance (Suppl. table 1). A third candidate region was identified on BTA15, containing two signal regions associated with two olfactory receptor genes (OR4A2C and OR4A2I).

#### Signatures of selection found in Simgud only

Several Simgud-specific genomic regions under potential selection were identified, notably on BTA2, BTA3, BTA4, BTA14, BTA20, and BTA22. The signal on BTA3 harboured NHLH2 and an uncharacterized gene, while on BTA20 the region under selection contains the RGS7BP and ADAMTS12 genes. The signal on BTA22 harbors the MITF gene. Moreover, these signatures of selection overlapped with multiple known QTLs linked to economically and biologically important traits (Suppl. table 1). For instance, the signal on BTA3, spanning 13.29–13.37 Mb harbors overlapping QTLs for milk protein percentage and shear force, indicating selection for both dairy production and meat quality traits. Likewise, the detected signal on BTA20 (14.84–14.86 Mb) includes a QTL associated with heel depth, which may relate to structural soundness and animal mobility. The most prominent cluster was observed on BTA22 (31.59–31.93 Mb), where numerous QTLs for white spotting, hoof and leg disorders, milk fat yield, bovine tuberculosis susceptibility, and shear force were co-localized. Moreover, GO analyses of genes in regions under selection in Simgud cattle (iHS) identified significant enrichment in taste-related pathways. Additional categories, includes G protein-coupled receptor signaling and chondrocyte differentiation (Suppl table 2). Consistently, the KEGG pathway analysis revealed Taste transduction as significantly enriched pathway (Suppl table 3 and Suppl. fig. S1).

#### Overlapping regions showing signatures of selection between Gudali and Simgud

Intra-population haplotype-based analyses using the iHS statistic revealed several fully or partially overlapping selection signals between Gudali and Simgud cattle (Table 1), highlighting common adaptive regions across chromosomes. The most prominent shared signature was observed on BTA6 (49–54 Mb), representing the longest and strongest signal in both populations. This region harbours the 5S ribomosal rRNA gene (5S_RNA). On BTA5 and BTA7, both populations exhibited selection signals involving genes related to sensory functions. In Gudali, the signal on BTA5 encompasses the OR6C8 gene, while in Simgud, an overlapping region harbored HMGA2 and two bitter taste receptor genes (TAS2R42 and TAS2R46), potentially influencing foraging behavior.

Likewise, Gudali exhibits a peak containing OR2O2 on BTA7 while Simgud showed broader overlapping signals involving multiple genes, including OR7A94, OR7H5P, and regulatory genes like ABLIM3 and AFAP1L1. On BTA24, a key shared region was identified encompassing the WDR7 gene in both Gudali and Simgud, with Gudali additionally showing selection on the neighboring TXNL1 gene. Overall, the shared signatures largely involve genes related to sensory perception, metabolism, and regulatory functions, indicating common selective pressures likely associated with environmental adaptation and feed-related behavior.

#### Tajima’s D Statistic Test

The Manhattan plots of Tajima′s D values are shown in 3. The means and standard deviations of the Tajima’s D statistic were 2.48 *±* 1.08 and 1.66 *±* 1.00 for Gudali and Simgud populations, respectively. A total of 12 and 17 non-overlapping selection signatures were observed in Gudali and Simgud, respectively (Table 2). The most striking cluster was located on BTA29, where contiguous windows spanning the first ≈0.5 Mb showed Tajima’s D values down to –2.09. In Simgud, the cluster of contiguous windows spans the first ≈0.6 Mb and exhibited Tajima’s D values as low as –2.63. Several of the identified regions overlap in both populations and contain genes with potential functional relevance, including CTNNA3 (BTA28) and GLDN on BTA10. Regions containing MSRB3 and LEMD3 on BTA5 were specific to Gudali, while COL5A1 (BTA11) was found only in Simgud.

**Table 2.**
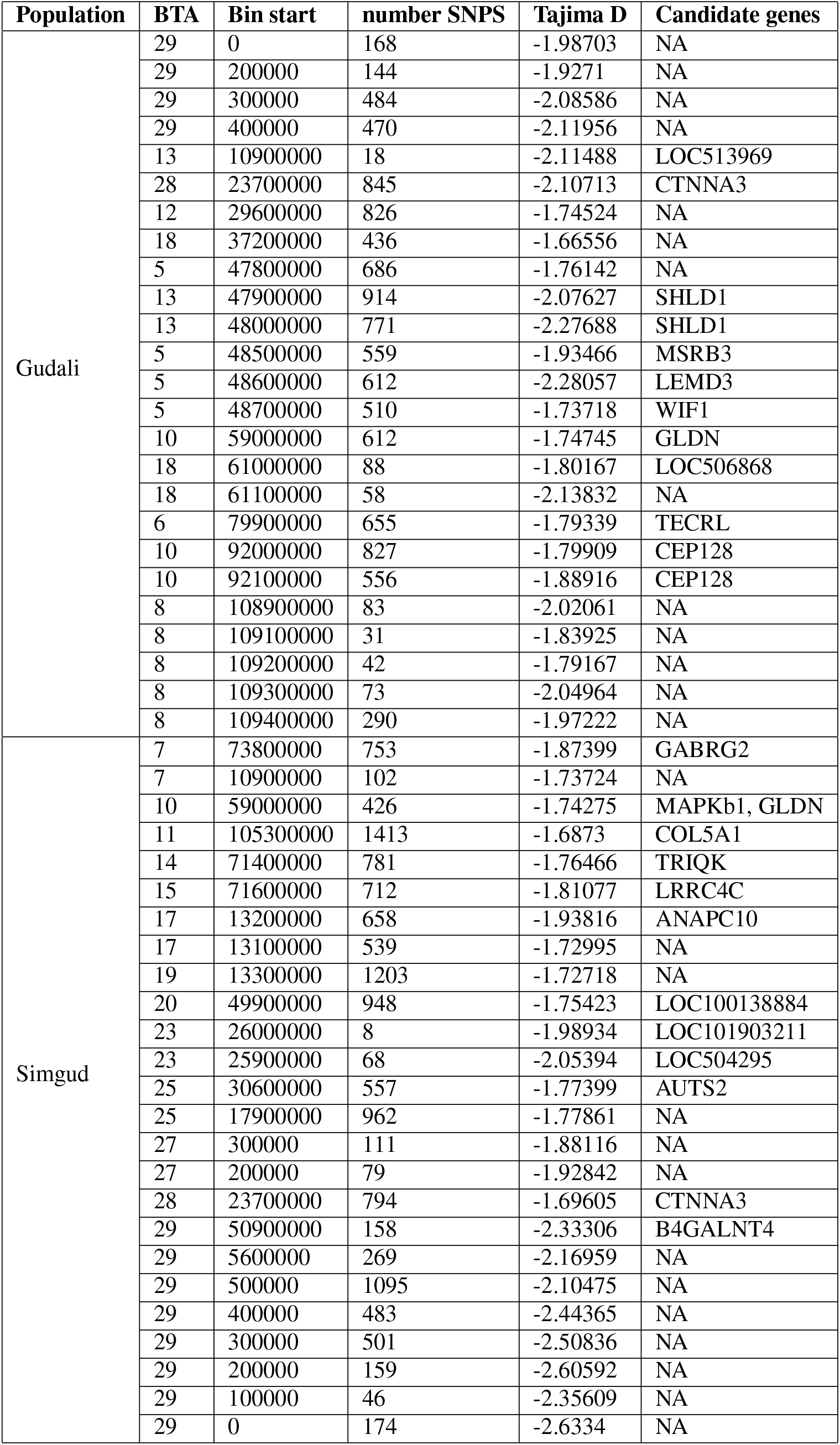
Candidate genes detected within signature of selection region using Tajima’s D statistic.

### Between population signals

#### Cross-population extended haplotype homozygosity (XP-EHH)

A cross-population extended haplotype homozygosity (XP-EHH) analysis was performed between Gudali and Simgud cattle to identify genomic regions under divergent selection. In total, 11 selection signature regions were detected and mapped to 12 genes (Fig. 3). In addition, the detected regions overlap with QTLs related to production, health, and fertility (Suppl. table 1). On BTA9, the signals were mapped to several genes such as C9H6orf203, BEND3, CD24 and LRP11. On BTA14, the QTL associated with dry matter intake lies within a region containing the EYA1 gene. On BTA16, overlapping QTLs for milk protein percentage and conception rate coincide with the TNR gene. On BTA20, within the region linked to milk protein percentage, the gene MTRR (methionine synthase reductase) is present. BTA23 harbours IRF4 and DUSP22 genes. On BTA29, a selection region associated with both bovine tuberculosis susceptibility and dry matter intake includes the GAB2 and TRNAC-ACA genes.

**Figure 3.**
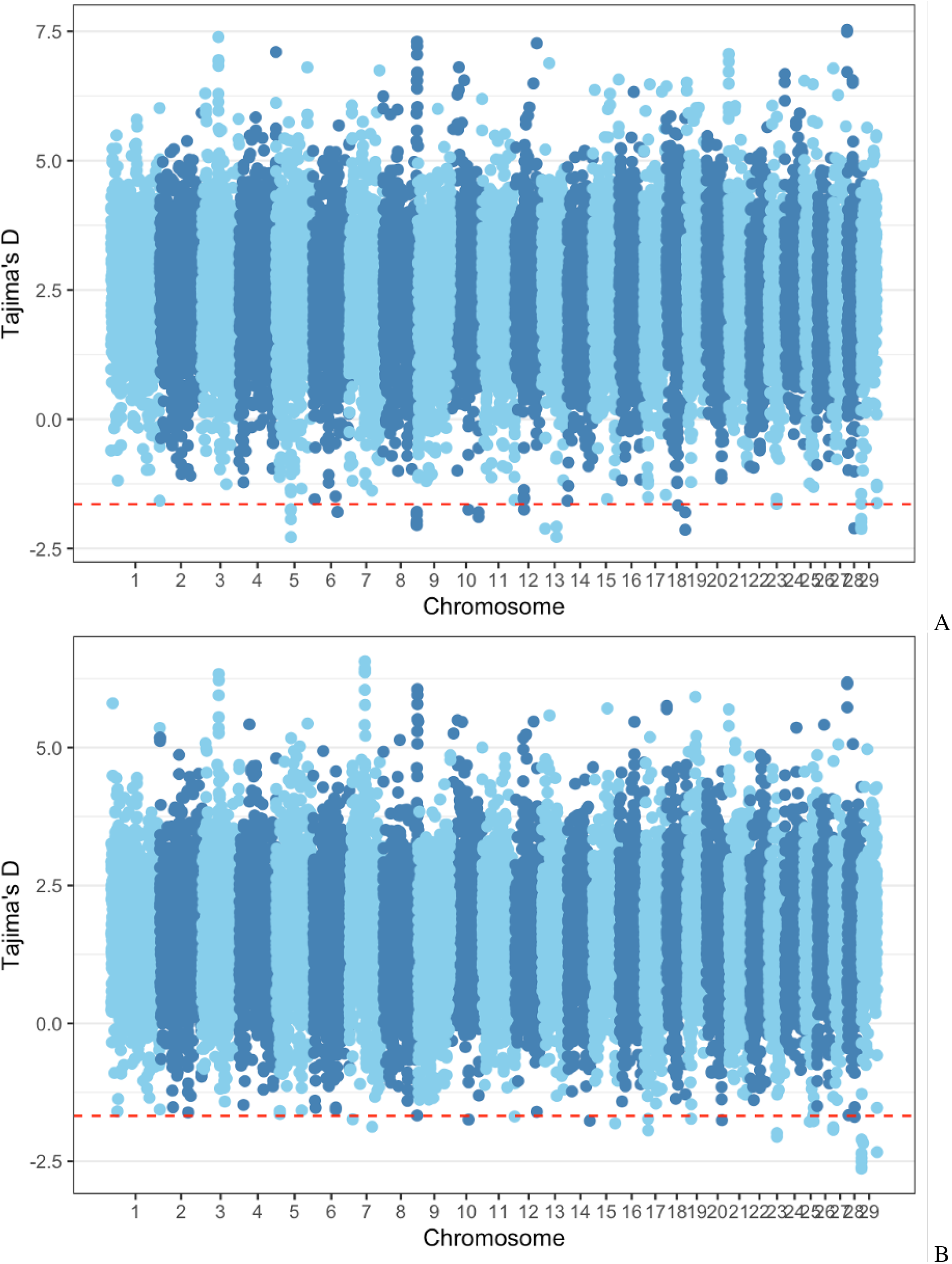
Manhattan plot of Tajima’s D values for indigenous Gudali (A) and crossbreed Simgud (B) cattle populations. The red dashed lines indicate the thresholds of −1.64, and −1.68 used to identify potential selection signatures in each population, respectively.

**Figure 4.**
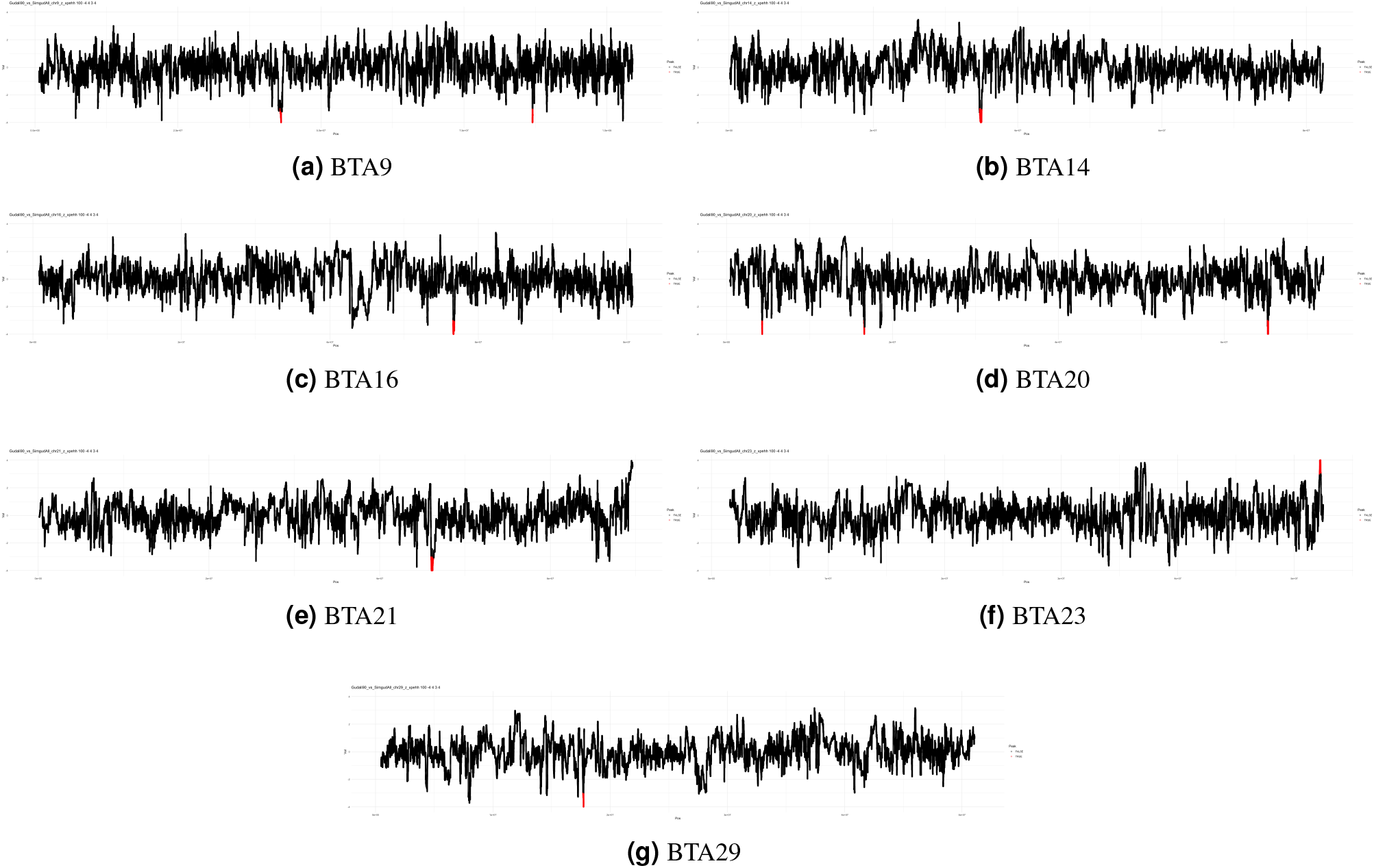
Cross-population standardized XP-EHH.

#### Population differentiation

By plotting the Z_F_ST_ for each 1,000 Kb genomic bin, a total of 98 candidate genomic regions differentiating Gudali and Simgud were detected on BTA2, BTA5, BTA7 and BTA12 (Figure 5). By investigating the gene content of these significant genomic bins, 55 genes with known function in cattle as well as in other species were identified (Suppl. table 4). For instance, differentiation analysis along BTA5 revealed a broad region of elevated genetic divergence spanning approximately 47.45 to 48.80 Mb, with F_ST_ values consistently exceeding 0.33 and peaking above 0.52. The upstream segment (47.45–47.90 Mb) showed moderate to high F_ST_ scores (0.33–0.44), highlighting genes such as GRIP1, HELB, IRAK3, TMBIM4, and HMGA2. A second, more pronounced differentiation peak was detected between 48.37 and 48.80 Mb, where the highest F_ST_ value (0.55) was recorded. This peak encompassed key genes including MSRB3, LEMD3, and WIF1. BTA7 showed a strong genetic differentiation candidate region (50–52 Mb) containing multiple windows with elevated F_ST_ values (0.32–0.36). This core region of 2 Mb on BTA7 is a candidate selective sweep or differentiation hotspot with CTNNA1, SIL1, MATR3, and NRG2 emerging as strong candidate genes. On BTA12, the peak region (29.05-29.7 Mb), contains three genes (FRY, RXFP2, MIR2299). GO enrichment analysis of genes under selection between Gudali and Simgud cattle identified serotonergic neuron axon guidance as the most significantly enriched process (Benjamini = 0.0026) (Suppl table 5). Additional enriched categories included cell adhesion, ascorbic acid transport, and response to toxic substances, while processes such as sodium ion transport and lung development showed moderate enrichment.

**Figure 5.**
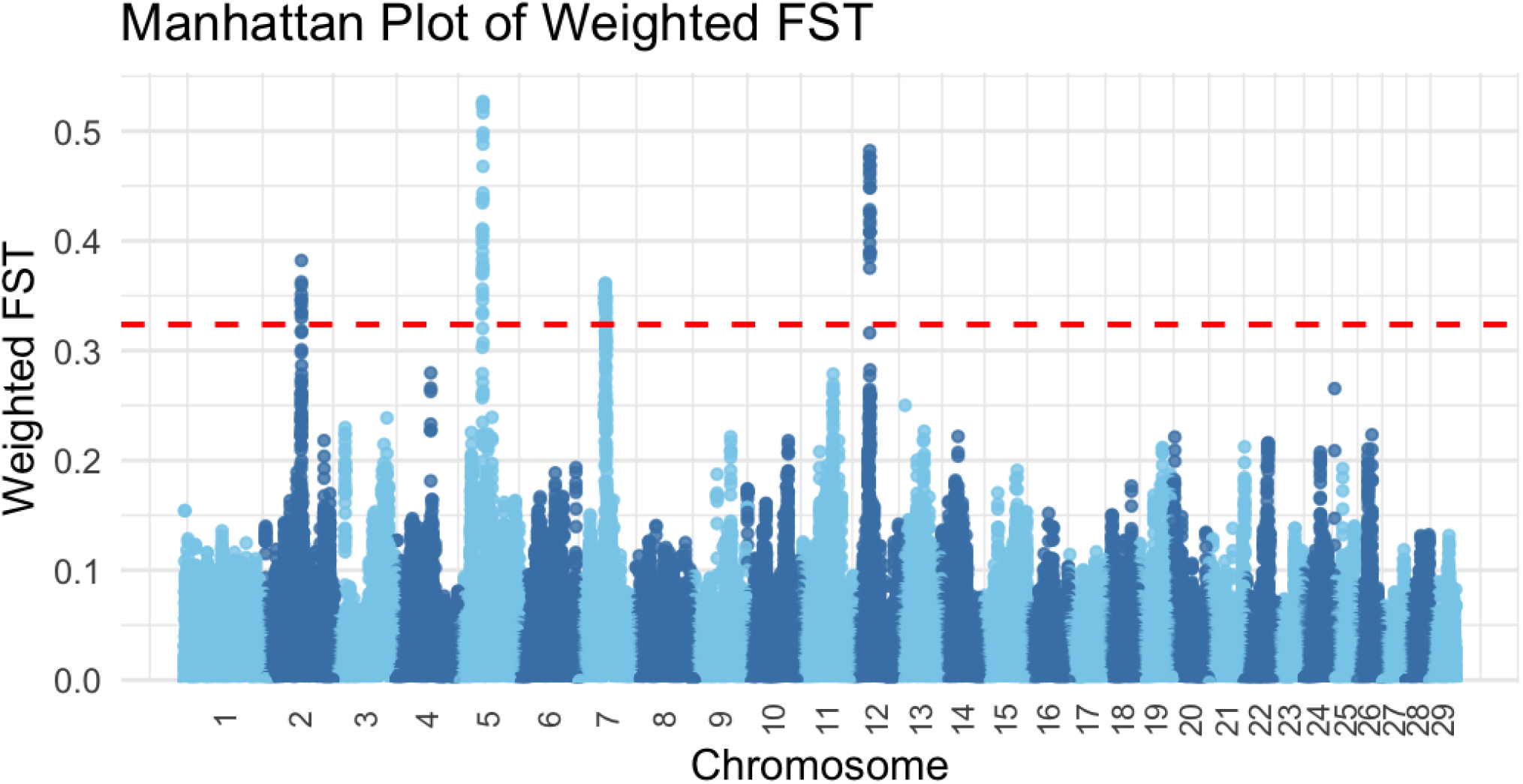
F_ST_-based differentiation between Gudali and Simgud populations.

### Tissue-specific candidate genes under selection

To investigate whether selection in Gudali and Simgud cattle acts on regulatory variation, we assessed whether the genes identified within the genomic regions under selection overlapped with cis-expression quantitative trait loci (cis-eQTL) from the CattleGTEx database. We used the RNA-seq data from 8,742 samples across 27 tissues, representing 27,607 expressed genes. Cis-eQTL mapping was performed per tissue, and tissue-specific significance was assessed using chi-squared and permutation-based tests. For the four genes detected in genomic region under selection in the cross-population analysis Table 4, gene enrichment analysis showed that testis has the strongest enrichment of cis-eQTL signals, consistent with its high transcriptional activity and complex regulatory landscape. Significant signals were also observed in Kidney and Lung, followed by more moderate enrichments in Oviduct, Lymph node, Spleen, and Adipose. In addition, selection in Simgud population, showed that selection may be acting through gene regulation, since strong tissue specific significance within Simgud was observed (table 5).

**Table 3.** Cross validation table of detected signals and annotated genes.

| BTA | Region start | Region end | Candidate genes |
| --- | --- | --- | --- |
| 14 | 34766569 | 35012294 | EYA1 |
| 16 | 56611544 | 56771917 | TNR |
| 20 | 4276182 | 4309319 | NA |
| 20 | 16601830 | 16659437 | NA |
| 20 | 65238206 | 65327978 | MTRR |
| 21 | 46045551 | 46119578 | BRMS1L, LOC101907487 |
| 21 | 46163787 | 46296002 | LOC107131617 |
| 23 | 52173712 | 52271299 | IRF4, DUSP22 |
| 29 | 17691538 | 17739396 | TRNAC-ACA, GAB2 |
| 9 | 42909961 | 43048057 | C9H6orf203, BEND3, LOC100137843, CD24, LOC614649 |
| 9 | 86968799 | 87037366 | LRP11 |

**Table 4.**
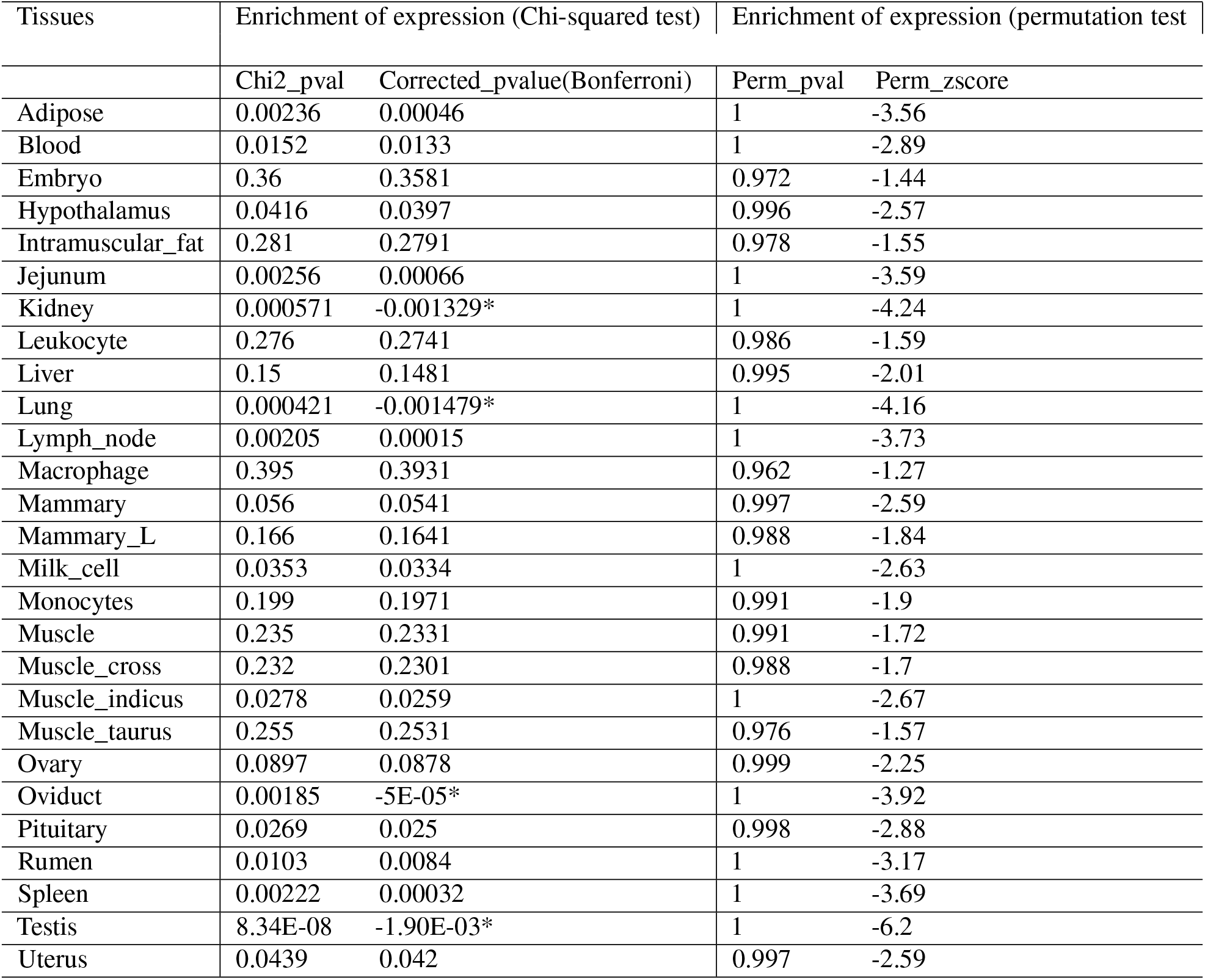
Enrichment of XP-EHH selection candidate genes in tissue-specific CattleGTEx over-expression database.

| Tissues | Enrichment of expression (Chi-squared test) |  | Enrichment of expression (permutation test) |  |
| --- | --- | --- | --- | --- |
|  | Chi2_pval | Corrected_pvalue(Bonferroni) | Perm_pval | Perm_zscore |
| Adipose | 0.00236 | 0.00046 | 1 | -3.56 |
| Blood | 0.0152 | 0.0133 | 1 | -2.89 |
| Embryo | 0.36 | 0.3581 | 0.972 | -1.44 |
| Hypothalamus | 0.0416 | 0.0397 | 0.996 | -2.57 |
| Intramuscular_fat | 0.281 | 0.2791 | 0.978 | -1.55 |
| Jejunum | 0.00256 | 0.00066 | 1 | -3.59 |
| Kidney | 0.000571 | -0.001329* | 1 | -4.24 |
| Leukocyte | 0.276 | 0.2741 | 0.986 | -1.59 |
| Liver | 0.15 | 0.1481 | 0.995 | -2.01 |
| Lung | 0.000421 | -0.001479* | 1 | -4.16 |
| Lymph_node | 0.00205 | 0.00015 | 1 | -3.73 |
| Macrophage | 0.395 | 0.3931 | 0.962 | -1.27 |
| Mammary | 0.056 | 0.0541 | 0.997 | -2.59 |
| Mammary_L | 0.166 | 0.1641 | 0.988 | -1.84 |
| Milk_cell | 0.0353 | 0.0334 | 1 | -2.63 |
| Monocytes | 0.199 | 0.1971 | 0.991 | -1.9 |
| Muscle | 0.235 | 0.2331 | 0.991 | -1.72 |
| Muscle_cross | 0.232 | 0.2301 | 0.988 | -1.7 |
| Muscle_indicus | 0.0278 | 0.0259 | 1 | -2.67 |
| Muscle_taurus | 0.255 | 0.2531 | 0.976 | -1.57 |
| Ovary | 0.0897 | 0.0878 | 0.999 | -2.25 |
| Oviduct | 0.00185 | -5E-05* | 1 | -3.92 |
| Pituitary | 0.0269 | 0.025 | 0.998 | -2.88 |
| Rumen | 0.0103 | 0.0084 | 1 | -3.17 |
| Spleen | 0.00222 | 0.00032 | 1 | -3.69 |
| Testis | 8.34E-08 | -1.90E-03* | 1 | -6.2 |
| Uterus | 0.0439 | 0.042 | 0.997 | -2.59 |

**Table 5.** Enrichment of candidate genes under selection in Simgud in tissue-specific CattleGTEx over-expression database.

| Tissues | Enrichment of expression (Chi-squared test) |  | Enrichment of expression (permutation test) |  |
| --- | --- | --- | --- | --- |
|  | Chi2_pval | Corrected_pvalue(Bonferroni) | Perm_pval | Perm_zscore |
| Testis | 4.72E-29 | 1.27E-27* | 1 | -11.48 |
| Lung | 3.01E-17 | 8.13E-16* | 1 | -8.76 |
| Hypothalamus | 4.51E-15 | 1.22E-13* | 1 | -8.3 |
| Pituitary | 7.02E-15 | 1.90E-13* | 1 | -8.22 |
| Kidney | 1.43E-14 | 3.86E-13* | 1 | -8.13 |
| Lymph_node | 1.20E-13 | 3.24E-12* | 1 | -7.77 |
| Spleen | 1.82E-13 | 4.91E-12* | 1 | -7.7 |
| Jejunum | 3.73E-13 | 1.01E-11* | 1 | -7.44 |
| Mammary | 3.90E-12 | 1.05E-10* | 1 | -7.16 |
| Ovary | 6.67E-12 | 1.80E-10* | 1 | -6.92 |
| Adipose | 1.10E-11 | 2.97E-10* | 1 | -7 |
| Rumen | 4.88E-10 | 1.32E-08* | 1 | -6.64 |
| Liver | 7.63E-10 | 2.06E-08* | 1 | -6.47 |
| Muscle_cross | 2.35E-09 | 6.35E-08* | 1 | -6.44 |
| Monocytes | 9.67E-09 | 2.61E-07* | 1 | -6.04 |
| Muscle | 4.38E-08 | 1.18E-06* | 1 | -5.77 |
| Muscle_indicus | 7.21E-08 | 1.95E-06* | 1 | -5.7 |
| Oviduct | 8.77E-08 | 2.37E-06* | 1 | -5.48 |
| Leukocyte | 1.77E-07 | 4.78E-06* | 1 | -5.52 |
| Intramuscular_fat | 2.08E-07 | 5.62E-06* | 1 | -5.39 |
| Uterus | 4.49E-07 | 1.21E-05* | 1 | -5.33 |
| Muscle_taurus | 1.06E-06 | 2.86E-05* | 1 | -5.07 |
| Milk_cell | 2.55E-06 | 6.89E-05* | 1 | -5.03 |
| Blood | 8.18E-06 | 2.21E-04* | 1 | -4.63 |
| Embryo | 1.53E-05 | 4.13E-04* | 1 | -4.51 |
| Macrophage | 0.000204 | 5.51E-03* | 1 | -3.9 |
| Mammary_L | 0.00907 | 2.45E-01 | 1 | -2.85 |

### Trait association of selection signatures

The detected signatures of selection were mapped to the CattleQTLdb database to identify overlaps between these signals and QTL. A total of 178 QTL intersect with the regions under selection as detected by iHS and XP-EHH analysis (Suppl. table 1). These are generally known quantitative trait loci (QTLs) associated with economically important traits, supporting their potential role in breed-specific adaptation and production performance.

## Discussion

This study investigates selection signatures in the local West African short horn zebu (Gudali) and its crossbreed with the Italian Simmental (Simgud), which was created some decades ago to improve beef production in Cameroon. We used several intra- and interpopulation test metrics and detected regions under selection. These are identifiable by their deviation from the expected occurrence in a neutral theory scenario^15^ and likely occurring in genomic regions harboring genes playing essential roles regarding the phenotypes under selection^16^. In a recurrent selection situation, some alleles are favoured over others, leading to an increase in their frequency in the population. This means they have a beneficial effect on individuals’ phenotypes, unlike in the neutral theory situation which assumes that genetic variants have no effect on the performance of individuals. Diversity in the region under selection is limited and there is a remarkably long-range haplotype^17^ around the markers favoured by selection. We considered the top 0.1% of the genomic windows identified as candidate selection signatures and merged adjacent windows to detect signals of potential recent selection. The crossbreeding initiative started only two decades ago: not enough time for chromosomal segments harbouring these signals to undergo disruption and recombination during meiosis. Genes found within the detected selection signatures highlight positive selection pressure related to adaptation. For both Gudali and Simgud, the mean Tajima’s D value observed across all genome regions was positive. This result was expected because breeding programs generally target several traits: in the case of the Gudali/Simgud crossbreeding initiative, traits such as adaptation, disease resistance and growth. Also, since only about 5% of the mammalian genome is influenced by natural selection^18^, the overall Tajima’s D value across the entire genome is expected to be positive (not strongly negative or zero). This positive value likely reflects the neutral or nearly neutral evolution dominating most of the genome, rather than strong selective sweeps that push Tajima’s D negative in the candidate regions we identified. To improve the robustness of our results, we have employed multiple methods since such approaches have been reported to improve the probability of detecting selection signatures with greater confidence^16^. However, it is worth mentioning that, because different methods for detecting signatures of selection rely on different statistical approaches, it is expected that they will generate different findings. By comparing and contrasting these, the methods can be used to complement each other^19^. Based on this, we have considered all of the identified signals, together with the candidate genes/QTL found within these signals. Mostly, genes co-localizing with signatures of positive selection were related to growth and development, fertility and reproduction, heat tolerance and environmental adaptation, immunity and disease resistance, production and quality traits.

### Growth and development

In Gudali, a strong selection signal was identified on BTA6 (4.9–5.3 Mb), the largest region under selection (456 Kb), which harbors the 5S ribosomal RNA gene. This region overlaps with multiple QTL for maturity rate, milk riboflavin and fatty acid content, and shear force. The high QTL density in this region indicates a genomic hotspot for developmental, milk composition, and meat quality traits. The 5S_RNA gene has also been detected under selection in North African cattle breeds^20^. The NHLH2 gene on BTA3 in Simgud, located near a QTL for marbling, is highly conserved and plays a role in body weight regulation^21^. In mice, its deletion leads to adult-onset obesity due to reduced voluntary activity without increased food intake^22^. This suggests a role in energy balance and physical activity, with implications for cattle growth and development. In both Gudali and Simgud, the GLDN gene found on BTA10 has previously been linked to skeletal development and metabolism in Sanjiang cattle^23^ and shows selection signatures in Mengshan cattle^24^. Its role in bone metabolism^25^ suggests an adaptive advantage for movement and survival in harsh landscapes. In Simgud, a fertility-growth hotspot on BTA5 (47.7–47.8 Mb) includes the HMGA2 gene, co-localizing with QTLs for fertility (first service conception, pregnancy rate, interval to first estrus, inseminations per conception), growth (body weight, stature), and endocrine traits (inhibin level). HMGA2 regulates PLAG1 and IGF2, both linked to growth and fertility traits^26,27^. It is associated with height in humans^28^, body size in animals^29,30^, and fertility in cattle^31,32^. HMGA2 was also found under selection in Gir and Holstein cattle^33,34^. The CTNNA1 gene on BTA7, detected in the differentiation analysis, has a role in muscle development. It has been associated with myostatin expression in skeletal muscle in Holstein-Friesian bulls^35^.

### Fertility and reproductive traits

On BTA1 (155.6–155.9 Mb), selection signals overlap QTL for milk composition, fertility (calving ease, conception, pregnancy rate), health, and longevity. Despite lacking an annotated gene, the diversity of overlapping QTL suggests selection on regulatory elements or linked loci, consistent with milk–fertility–health tradeoffs in dairy breeding^36^. Also on BTA1, another region (115.5–115.8 Mb) overlaps with QTLs for milk and protein yield, and methane production, suggesting selection for production efficiency. On BTA24 (55.94–55.98 Mb), TXNL1 and WDR7 were identified under selection. TXNL1 is a redox-active thioredoxin involved in oxidative stress protection^37^ and was found active in the corpus luteum during early pregnancy in Finnsheep^38^. WDR7 is involved in vesicle trafficking and neuronal communication^39,40^. The ADAMTS12 gene on BTA20 has been linked to pregnancy loss in Nellore cattle^41^, milk production and carcass weight^42,43^, and spontaneous preterm birth in humans^44^. It also influences morphogenesis during embryonic development^45^. The EYA1 gene on BTA14 overlaps with a QTL for dry matter intake and has been suggested to play roles in urinary tract development and physiological adaptation to drought or heat^46^. The RXFP2 gene on BTA12, identified via differentiation analysis, has pleiotropic effects on inguinoscrotal testis descent^47^, horn size in bighorn sheep^48^, and domestic sheep^49,50^. Its thermoregulatory role via nasal heat exchange has been proposed^51^.

### Heat tolerance and environmental adaptation

The DSCAM gene on BTA1 was identified as being under strong selection in Simgud and is involved in neural development and immune regulation. It contributes to neural circuit formation and modulates immune responses^52–54^, making it relevant to both behavior and disease resistance. MSRB3 on BTA5 showed evidence of selection in both iHS and F_ST_ analyses and is linked to oxidative stress resistance and heat tolerance^55^. It also influences auditory traits, body size, and growth in several species^29,56–62^. It has been previously reported in Ethiopian cattle^63^, Tibetan dogs, and sheep^64,65^. TAS2R42 and TAS2R46, located near HMGA2, encode bitter taste receptors. These help cattle avoid toxic plants^66,67^, suggesting dietary adaptation in grazing systems. LEMD3 and WIF1, also on BTA5, are related to melanogenesis and coat color, contributing to heat tolerance via pigmentation adaptations^68,69^.

### Immunity and disease resistance

OR genes such as OR6C8, OR2O2, OR4A2I, and OR4A2C were enriched in selection regions on BTA5 and BTA15. OR genes, the largest gene family in vertebrates, are critical for foraging, social behavior, and parasite avoidance in tropical environments^20,70–75^. The GAB2 gene on BTA29 overlaps with QTLs for tuberculosis susceptibility and dry matter intake. It has been identified under selection in trypanosome-resistant African breeds^76^. IRAK3, GRIP1, and HELB on BTA5 are all immunity-related genes. IRAK3 modulates inflammatory responses and has been found under selection in several African cattle breeds^6,77–79^. GRIP1 supports glucocorticoid anti-inflammatory effects^80^. HELB is involved in DNA repair and replication stress response^81^. MITF on BTA22 is involved in pigmentation and immunity, linked to QTLs for tuberculosis susceptibility, white spotting, and hoof and leg disorders^82–85^. SIL1 on BTA7 is involved in protein folding and was detected in ROH islands in Chinese breeds^86^.

### Production, health, and quality traits

The region on BTA5 (57.67–57.68 Mb) harbors QTLs for milk fat yield, coat color, somatic cell count, and sole ulcer susceptibility. This convergence indicates co-selection on productivity and robustness traits^87–90^. The dense QTL cluster on BTA22 (31.59–31.93 Mb) includes traits related to milk fat yield, shear force, tuberculosis, and conformation (e.g., hoof disorders, white spotting), and houses the MITF gene. This region likely reflects pleiotropic selection for health and production. COL5A1 on BTA11 in Simgud, a collagen gene associated with joint laxity and connective tissue disorders in humans and animals^91–94^, is linked to mobility and robustness in grazing systems. CTNNA3 on BTA28, found in overlapping selection regions of both populations, plays a role in mammary gland development and milk production^95^.

## Functional analysis

The enrichment of neuronal signaling and antioxidant metabolism pathways in F_ST_ analysis suggests that genetic divergence between Gudali and Simgud cattle involves behavioral adaptation and oxidative stress tolerance. In fact, serotonin influences a wide range of physiological and behavioral processes in animals^96^. Moreover, the serotonergic system encompasses embryonic development, pulmonary, vascular, cardiac, gastrointestinal, and reproductive functions^97^. These results highlight that selection in these populations extends beyond production traits to encompass adaptive processes important for survival and robustness in low-input systems.

The enrichment of taste-related pathways, including both GO terms and the KEGG Taste transduction pathway, indicates that selection in Simgud cattle may influence dietary adaptation and feed preference, potentially affecting foraging behavior and nutrient acquisition in their grazing environment. This result correlates with our findings about bitter taste receptor genes (TAS2R42 and TAS2R46), identified as being under selection in Simgud, thus theoretically assisting the hybrid animals to avoid grazing on toxic plants^66,67^. This result further reinforces the enrichment of olafctory genes found (OR6C8, OR2O2, OR4A2I, and OR4A2C) in selection regions on BTA5 and BTA15. Selection in these regions may be considered as an adaptation mechanism for foraging, social behavior, and parasite avoidance in tropical environments^20,70–75^. The involvement of GPCR signaling pathways reinforces the role of sensory signal transduction in adaptation, while enrichment of chondrocyte differentiation suggests selective pressure on skeletal growth and morphology, reflecting environmental or management-related adaptations. Together, these results highlight that both sensory perception and growth-related processes contribute to the adaptive profile of Simgud cattle.

### Tissue specificity of candidate genes under selection

The tissue-specific enrichment of cis-eQTLs in genomic regions under selection suggests that selection may be acting through gene regulation, particularly in Testis, Kidney, and Lung. This is biologically plausible given the known roles of these tissues in reproduction, metabolic homeostasis, and environmental adaptation—key traits in tropical and crossbred cattle. The strong signal in Testis aligns with previous findings about its well-documented high transcriptional activity and unique regulatory landscape^98–100^. Meanwhile, kidney and lung are central to thermoregulation, water balance, and oxygen exchange, which are crucial in heat-stressed or resource-limited environments. This results support the hypothesis that cis-regulatory variants contribute to local adaptation and productivity in cattle^101^. Even after permutation analyses, although p-values were uniformly close to 1, large negative permutation Z-scores (up to –6.2 in Testis), highlight consistent biological signal in those tissues (notably Testis, Kidney, and Lung) which retained robust significance after multiple testing. This result highlights these tissues as priority targets for follow-up studies of gene regulation in livestock.

Tissue-specific enrichment of cis-eQTL was observed exclusively in the Simgud crossbred population, with no significant signal detected in the indigenous Gudali breed. This difference may reflect increased regulatory activity in crossbreds arising from the combination of divergent parental alleles and exposure to stressful environmental conditions. Notably, higher gene expression levels have been reported in *Bos taurus × Bos indicus* hybrids when the sire was of taurine origin, a pattern likely linked to reduced tolerance to stressors such as heat and drought^102^. The Simmental ancestry likely contributes novel cis-regulatory variants, enhancing the detectability of tissue-specific expression patterns, particularly in metabolically active tissues like testis, kidney, and lung. In contrast, the Gudali breed may exhibit reduced regulatory variability due to long-term local adaptation and stabilizing selection, which can constrain the expression landscape and diminish observable eQTL enrichment in public datasets. Additionally, current eQTL databases such as CattleGTEx are taurine-centric^11^, potentially under-representing regulatory variants unique to African indigenous breeds. These findings highlight the importance of considering regulatory variation in crossbreeding programs, as introduced cis-eQTL may influence traits related to fertility, metabolism, and disease resistance. However, care must be taken to preserve the finely tuned regulatory architecture of locally adapted breeds like Gudali, whose reduced expression variability may reflect evolutionary optimization to tropical environments. Future selection strategies should therefore balance introduced genetic diversity with the conservation of indigenous regulatory adaptations that underpin resilience and productivity under local conditions.

However, permutation analyses revealed that these signals may simply result from not correcting for background biases in enrichment analysis. In fact, traditional enrichment tests based on contingency tables, such as chi-square or hypergeometric tests, do not correct for background biases including gene length, gene expression level, or gene-gene dependencies^103^.

## Conclusion

In this study, we report for the first time the identification of candidate regions for signatures of positive selection in the genome of an indigenous indicine cattle (Gudali) of Cameroon and its crossbreed with Euopean taurine (Simgud). We applied a comprehensive suite of genomic tools — including iHS, Tajima′s D, XP-EHH, F_ST_, and eQTL mapping — the convergence of selection signals across which highlights robust candidate genes. These genes may contribute to important adaptive traits including thermotolerance, food scarcity, immunity, and productivity in tropical environments. The integration of expression data through eQTL analysis further strengthens the functional relevance of these loci, linking genomic variation to gene regulation. These findings enhance our understanding of local adaptation and genetic diversity in African cattle and identify valuable targets for future genomic selection and breeding programs tailored to sustainable livestock production in challenging environments.

## Methods

### Animal material and imputation

The animal material and the data collection process, as well as the two-step imputation of 100K SNP to HD SNP then whole genome, is explained in Matenchi & Hegarty (2025)^7^.

### Selection signatures

We performed haplotype-based analysis to detect potential selection signature regions that may arise from continuous use of artificial selection^16^. We used the haplotype-based integrated haplotype score (iHS) test^104^ to investigate potential regions under recent selection in the Gudali and Simgud populations, and we contrasted the two breeds with the cross-population extended haplotype homozygosity (XP-EHH) test^17^. Additionally, we used the Tajima’s D^105^ and F_ST_ statistics for further genetic diversity analysis.

#### Within population

##### integrated Haplotype Scoring (iHS)

The SNP-wise iHS analysis for each breed was performed using the hapbin program^106^. We specified parameters to exclude variants with minor allele frequency greater than 10% (–*minmaf 0*.*01*) and that the integral of the observed decay of EHH (i.e. iHH) should be calculated until a cutoff of 0.1 (–*cutoff 0*.*1*)^104^. The integrated EHH - denoted iHH0 and iHH1 when computed for the ancestral and the derived allele respectively - was used to calculate the unstandardised iHS and the result standardised to have a mean of 0 and a variance of 1 following Voight et al. (2006)^104^. The SNP-wise iHS statistic (Z-score) identified peaks were called using an in-house R script, detecting genomic regions in windows of 100 Kb, in which Z-score (standard iHS) exceeds a defined maximum threshold before dropping below the defined minimum on both sides. The thresholds were set to 3 and 2 for the maximum and minimum respectively following Dutta et al. (2020)^107^. Signals in detected genomic regions were retrieved from the Ensembl repository using biomaRt and mapped to the Ensembl ARS-UCD1.2 build version 96 and scanned for overlaps to detect potential candidate genes under selection.

##### Tajima’s D Test Statistic

Another metric used for selection signature detection is the Tajima’s D statistic, which compares the the number of segregating sites (S) and the nucleotide diversity, which is the average number of pairwise nucleotide differences between sequences in a sample^105^. By comparing these two measures of genetic diversity, the statistic aims to distinguish between a DNA sequence evolving randomly and one evolving under a nonrandom process, such as selection, demographic expansion, contraction, or introgression. Under evolutionary neutral conditions, its value is expected to be zero. Deviations from this neutral model indicate either fixation of alleles or rare alleles (negative Tajima’s D values) or balancing selection, which reflects an abundance of intermediate allele frequencies (for positive Tajima’s D values)^105,108^. The Tajima’s D statistic was computed in windows of 100 Kb, considering only windows with at least 5 markers. A stringent cutoff was applied, with only the bottommost 0.1%, windows retrieved and adjacent selection signatures merged.

#### Between population

##### Cross-Population Extended Haplotype Homozygosity (XP-EHH)

Phenotypic variability between cattle breeds, as shaped by genetic adaptation, was assessed using the cross-population extended haplotype homozygosity statistic (XP-EHH). This statistic considers that strong selection in an allele results in a faster accumulation of that allele frequency in the genome than recombination can break down the haplotype on which it resides. Based on this statistic, a high haplotype homozygosity is expected in one population compared to another, if they are under differential selective pressures^17^. Peak calling was performed following Dutta et al. (2020)^107^. XP-EHH scores were smoothed by averaging across 1000 SNP windows and putative selective sweep regions were those with an absolute standardised XP-EHH >4. The start and end coordinates of these regions were defined to be where the XP-EHH scores fell back below 3. The Ensembl ARS-UCD1.2 served as a reference to detect candidate genes that may be under selection, using BiomaRt.

##### Genetic differentiation

We performed a genome-wide scan for potential signatures of selection by calculating pairwise weighted F_ST_ values across sliding non-overlapping 100 Kb windows with a 25 Kb step size in BEDTools v 2.28.0^109^. To visualize the distribution of genetic differentiation, a Manhattan plot was constructed. A genome-wide significance threshold was set at the 99.9th percentile of the weighted F_ST_ distribution, whereby only the top 0.1% of windows were considered as candidate regions under putative positive selection. This threshold was indicated by a horizontal red dashed line on the plot. All computations and visualizations were performed in R using the dplyr and ggplot2 packages.

### Bioinformatic analysis

Candidate genes within significant regions were annotated based on the bovine reference genome. Functional enrichment analysis was performed using DAVID to identify overrepresented Gene Ontology (GO) biological processes and KEGG pathways among the candidate genes. Significance was assessed using P-values adjusted for multiple testing (Benjamini-Hochberg), and fold enrichment was calculated to evaluate the magnitude of overrepresentation. This approach allowed us to link genomic regions under selection to biological functions and adaptive traits relevant to breed-specific performance and environmental resilience.

### Identification of tissue-specific candidate genes under selection

For the detection of tissue-specificity of candidate genes detected under selection from previous sections, we performed a cis-eQTL analysis using the gene expression data from CattleGTEx^11^, specifically the dataset of Fang *et al*.^110^. This dataset consists of 8,742 RNA-seq samples, 27,607 genes across 27 different tissue (adipose, blood, embryo, hypothalamus, intramuscular_fat, jejunum, kidney, leukocyte, liver, lung, lymph_node, macrophage, mammary, mammary_lactating, milk_cell, monocytes, muscle, muscle_cross, muscle_indicus, muscle_taurus, ovary, oviduct, pituitary, rumen, spleen, testis, uterus). Candidate genes were derived from cross-validation signatures of Gudali and Simgud cattle and mapped to Ensembl gene IDs using the bovine GTF annotation.

Cis-eQTL summary statistics (nominal and permutation-based) were processed per tissue. For nominal files, genes with p-values < 0.05 were considered significant; for permutation-based files, genes with false discovery rate (FDR) < 0.05 were considered significant. Significant genes were then compared to the candidate gene list to identify overlaps.

To ensure an unbiased enrichment framework, we defined the set of all genes tested in CattleGTEx by merging unique gene_ids across tissues (all_genes_union) and also retained per-tissue tested gene sets. This allowed candidate overlaps to be evaluated against the correct background of genes actually tested in each tissue. Enrichment of selection candidate genes among significant eGenes was then assessed using two complementary approaches: (i) a Chi-squared test for overrepresentation against the background set, and (ii) a permutation test, where candidate gene labels were randomly shuffled across the tested gene sets to compute empirical p-values and Z-scores. To visualize tissue-specific significance of candidate genes, a binary matrix was generated where rows represented candidate genes and columns represented tissues; entries indicated whether a gene was significant in a given tissue.

All analyses were performed in R using the data.table, and GenomicRanges, ensuring reproducibility and scalability across multiple tissues. Enrichment tests were considered significant at p < 0.05 (Chi-squared) or with permutation-derived empirical p < 0.05 and Z > 1.96.

### Trait-association of selection signatures

To investigate the potential functional relevance of the identified selection signatures, we mapped them to known quantitative trait loci (QTL) using data from the Cattle QTL Database (https://www.animalgenome.org/cgi-bin/QTLdb/BT/index), which contains a total of 196,904 QTL and phenotype association records. Genomic coordinates of the selection signatures were checked against those of the QTL using the intersect function in BEDTools^109^, allowing identification of overlapping regions associated with economically and biologically important traits in cattle.

## Acknowledgements

We extend our sincere gratitude to the National Livestock Company of Cameroon (SODEPA) and its General Director, Mr. Koulagna Koutou, for initiating and supporting this study. We also thank Dr. James Prendergast for his valuable assistance in editing the scripts used in the analysis.

## Author contributions statement

YPM conceived the experiment(s), performed analysis and wrote the initial draft, MH supervised the work and assisted with data formatting. All authors contributed to writing the final submitted manuscript.

