## Supplementary figures and images for "Genomic and regulatory basis of adaptation in Cameroonian Gudali and Simgud Cattle"

### Supplemental figure S1

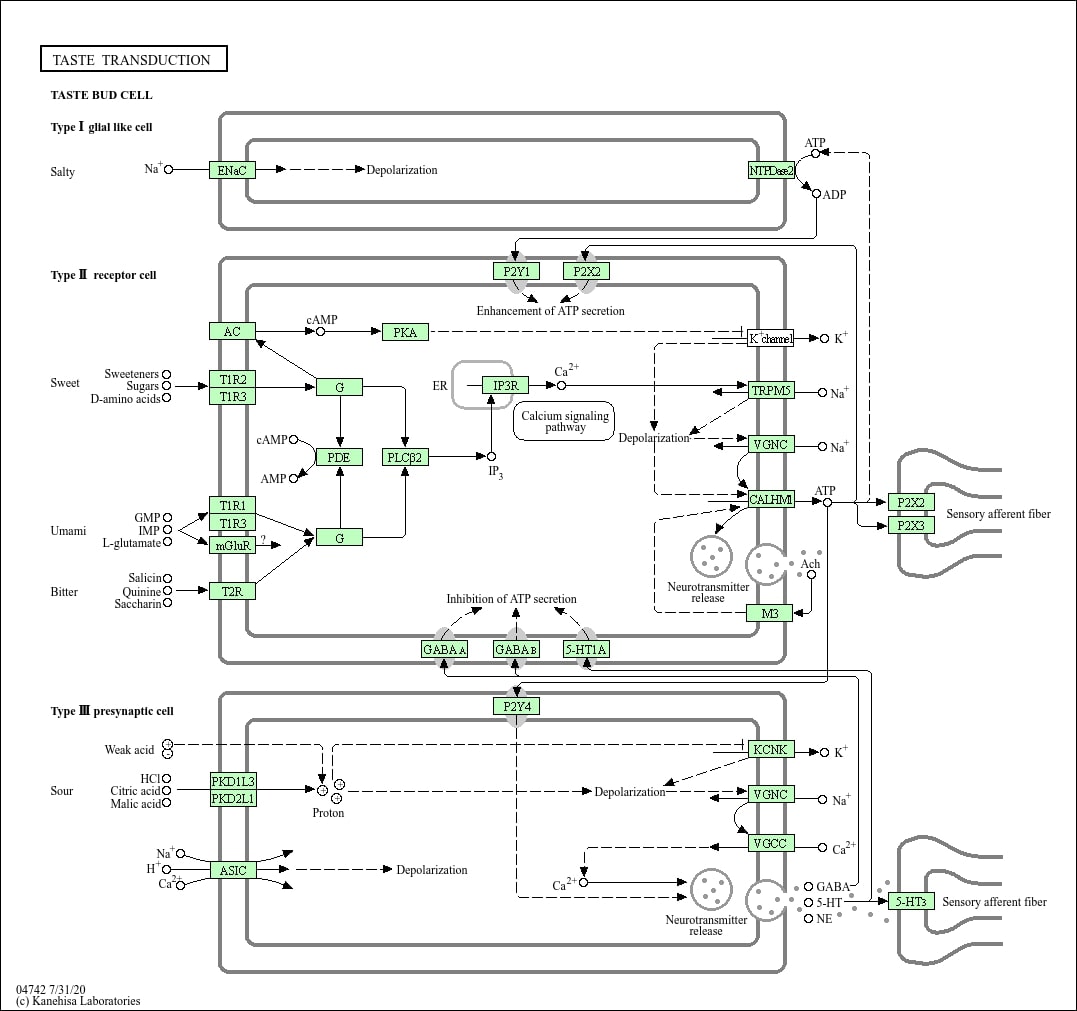
